# A Bioinformatic Pipeline for Consensus Taxonomic Classification of Long-Read Amplicons

**DOI:** 10.64898/2026.04.29.721641

**Authors:** Ashley A. Paulsen, Breah LaSarre, Drew Delp, Gwyn A. Beattie, Larry J. Halverson

## Abstract

Characterizing community composition is fundamental to understanding microbial community function. Recent advances in Oxford Nanopore Technology (ONT) long-read sequencing now allow community profiling using full-length gene amplicons, affording better taxonomic resolution than standard short-amplicon Illumina sequencing. However, robust ONT-compatible profiling workflows are lacking. To address this, we have created the Amplicon Consensus Taxonomy (ACT) pipeline for classifying long-read amplicons. ACT combines output from three existing pipelines – Emu, Sintax, and LACA – to leverage the strengths of each while offsetting their individual limitations. We also developed the ACT database (ACT-DB), a sequence-similarity-aware reference database that clusters highly similar sequences into multi-taxa groups to reduce overclassification. We benchmarked ACT performance against Emu and Sintax using a defined simple mock community, simulated datasets, and a complex rhizosphere community supplemented with novel species. While ACT exhibited generally comparable or superior performance across datasets, ACT demonstrated a marked advantage over Emu and Sintax in identifying novel and low-abundance taxa in both simple and complex communities, resulting in significantly higher species-richness estimates that better reflected those observed in prior Illumina amplicon studies. Furthermore, by clustering ambiguous reference sequences, ACT-DB allowed ACT to resolve reads to meaningful multi-species groups, improving resolution without coercing artificial precision. Together, ACT and ACT-DB form a robust long-read amplicon profiling workflow that confidently identifies known species while reducing overclassification and preserving low-abundance and unknown taxa.

**IMPORTANCE:** Microbial communities are frequently characterized by amplicon sequencing of marker genes, such as the bacterial 16S rRNA gene and fungal ITS region. Historically, the standard profiling method has been Illumina sequencing of 200-300 bp amplicons, but improved accuracy of ONT long-read sequencing means it is now possible to sequence amplicons spanning full genes of any size, prompting the need for tools optimized for long amplicons. Here, we describe the ACT bioinformatic pipeline for assigning taxonomy to amplicons of any length. We evaluated ACT performance using full-length 16S amplicon data relative to that of two commonly used pipelines. Additionally, we developed a sequence ambiguity-aware ACT database (ACT-DB) of 16S rRNA sequences to further improve classification accuracy and resolution.

## INTRODUCTION

A fundamental step in understanding how microbial communities contribute to ecosystem processes is characterizing community composition. Historically, prokaryotic community profiling has relied on Illumina-based sequencing of 16S rRNA gene amplicons (1–5). However, short Illumina amplicons capture less than 30% of the full 16S rRNA gene and cannot discriminate many taxa beyond the genus level (6, 7). Similar limitations exist for profiling microbial eukaryotes using the 18S rRNA gene or Internal Transcribed Spacer region. Consequently, there is widespread interest in high-throughput methods using longer amplicons to improve taxonomic resolution and facilitate deeper ecological insights.

Oxford Nanopore Technology (ONT) long-read sequencing has emerged as a promising cost-competitive approach for higher-resolution community profiling using full-length marker genes. With improved flow-cell chemistry and base calling models, ONT sequencing accuracy (>99%) is now sufficient for amplicon profiling. However, translating full-length marker gene data into reliable community profiles remains challenging because ONT-compatible workflows are limited in capability. Illumina-based pipelines are designed to cluster sequences <500 bp and require uniform amplicon length; however, full-length amplicons of 16S, 18S, and ITS marker regions vary in length across taxa. The few algorithms that can cluster variable length amplicons are computationally intensive with comparatively poor accuracy for long reads (8). Hence, community profiling using ONT reads requires alternative pipelines.

Several open-source tools for ONT amplicon profiling are available, each with distinct strengths and limitations. One popular tool is Emu (9), which has been incorporated into multiple workflows including EasyAmplicon2 (10), TRANA [https://github.com/genomic-medicine-sweden/TRANA], RESCUE (11) and Cosmos-Hub (cosmos-hub.com). Because of its probabilistic framework and error profile optimization, Emu excels at species-level classification of known taxa. Specifically, Emu uses an expectation-maximization algorithm to account for lower accuracy in long reads and iteratively refines abundance estimates based on read alignments to reference sequences. However, if Emu encounters a read from an unknown species, it will either overclassify the read as the closest known species or exclude the read as unclassified (7, 9). Thus, Emu is poor at classifying unknown species, which include both novel species that have not yet been sequenced and sequenced species that are absent from the reference database. Another limitation is that Emu can fail to detect low-abundance species, either because the species abundance is below the detection limit or because a rare species shares sequence similarity with dominant community members. For example, if Emu identifies 99% of reads as species A and 1% as closely related species B, Emu may incorrectly attribute the 1% to sequencing error and misclassify species B reads as species A. These limitations constrain Emu’s effectiveness for profiling microbial communities that are rich in rare and unknown taxa, like those in soil, plant, gut, and marine environments.

An alternative tool that can detect rare and unknown taxa is Sintax (12). Like Emu, Sintax assigns taxonomy using a reference database. Unlike Emu, Sintax provides bootstrap confidence estimates for each read at each taxonomic rank. By applying user-defined confidence thresholds, Sintax can assign read taxonomy at higher ranks even if species cannot be assigned. However, Sintax’s k-mer-based classification strategy is sensitive to sequencing errors. Consequently, Sintax detects rare and novel species but at the risk of misclassifying low-quality reads and at the expense of reduced classification confidence for known taxa.

Another limitation is that Sintax collapses distinct unknown species from the same genus into a single genus-level taxon and thus does not capture intra-genus diversity among unknown species. This diversity could be captured using similarity-based clustering of amplicons into Operational Taxonomic Units (OTUs), as is done for Illumina reads; however, algorithms for clustering long-reads have generally had performance issues, as noted above. Fortunately, the new Long Amplicon Consensus Analysis (LACA) workflow (13) addresses this challenge by integrating multiple long-read clustering tools to generate consensus OTUs with substantially lower error rates relative to the individual tools. The consensus OTUs can then be assigned taxonomy by alignment with reference databases. Despite potential limitations in detecting rare taxa, LACA reproducibly identifies genus- and species-level OTUs for community members with relative abundances >0.1%, illustrating the utility of LACA for profiling complex communities (13).

Here, we introduce the Amplicon Consensus Taxonomy (ACT) pipeline, which reconciles outputs from three tools – Emu, Sintax, and LACA – to generate consensus taxonomic assignments at the individual read level. We also introduce a sequence-similarity-aware reference database, the ACT database (ACT-DB). Unlike standard databases, the ACT-DB clusters highly similar sequences into multi-taxa groups on a per-gene-copy basis. This approach reduces misclassification when 16S rRNA sequences cannot reliably distinguish among closely related taxa while enabling species-level classification for other 16S rRNA copies having adequate resolution. We validated the performance of the ACT pipeline using full-length 16S rRNA ONT data from both a simple mock community and a complex soil community, further evaluated it using *in silico* benchmarking, and compared it with the Emu or Sintax pipelines using four standard reference databases and the ACT-DB. Our results demonstrate that the ACT pipeline enhances classification accuracy and diversity estimation by cross-validating taxonomic calls and incorporating clustering functions. These capabilities are further strengthened when coupled with the ACT-DB, which mitigates classification errors arising from biological sequence ambiguity. Together, the ACT pipeline and the ACT-DB provide a foundation for robust, high-resolution profiling of microbial communities using long-read amplicons.

## METHODS

### Amplicon consensus taxonomy (ACT) workflow

An overview of the ACT workflow is illustrated in Fig. 1. The ACT pipeline (hereafter ACT) accepts sequencing reads in fastq or fastq.gz format. If input data are multiplexed reads, ACT performs pre-processing that includes i) minimal length and quality filtering using chopper (14) to remove extreme outliers, with an upper length threshold greater than twice the expected amplicon size to preserve concatenated reads (11) that may be split during demultiplexing, and ii) demultiplexing and trimming with Cutadapt (15). Demultiplexing parameters allow users to specify barcode sequences, placement, and error tolerance. Following pre-processing, or for reads already demultiplexed and trimmed, ACT performs sample-specific filtering of sequences to a user-specified length range based on expected amplicon size, accommodating libraries with mixed amplicon lengths. ACT documentation includes a NanoPlot (14) helper script to inform filtering settings.

**Figure 1.**
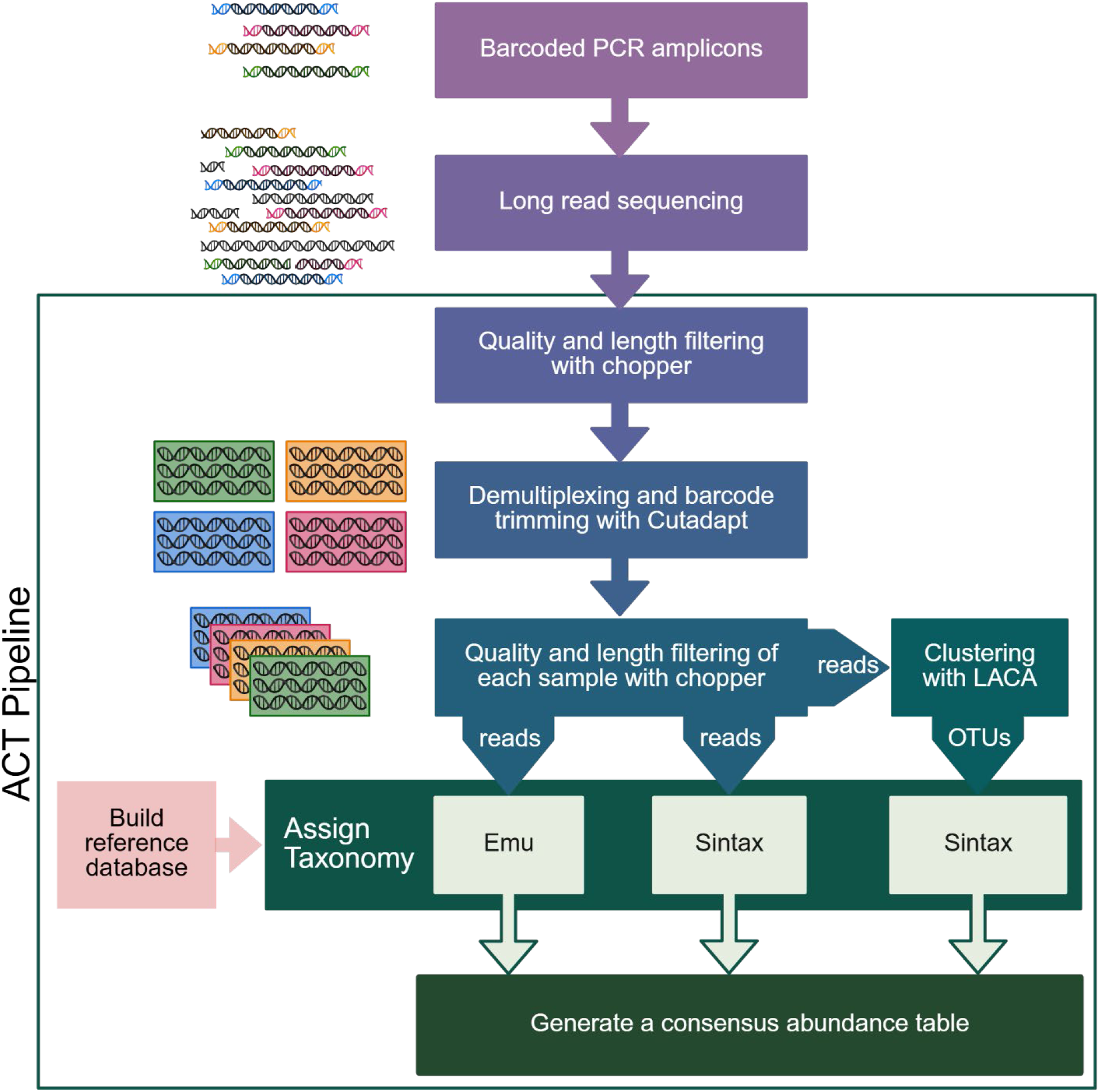
Amplicon Consensus Taxonomy (ACT) pipeline overview. Pipeline components are within the box while those above the box show steps for generating the data. Arrows that are labeled “reads” represent an identical set of reads provided to the Emu, Sintax, and Long Amplicon Consensus Analysis (LACA) pipelines. The arrow labeled OTUs portrays the representative sequence for each operational taxonomic unit designated by LACA for assigning taxonomy by Sintax. Created in https://BioRender.com

Filtered reads are then clustered and assigned taxonomy, which are performed simultaneously for efficiency. Emu and Sintax independently assign taxonomy to each read, while LACA clusters all reads into OTUs and generates a read ID-to-OTU map and a fasta file of OTU consensus sequences. The OTU consensus sequences are then assigned taxonomy by Sintax. The Emu output includes both the original un-polished Minimap2 alignments and the final expectation-maximization polished results. Emu is not able to assign taxonomy to OTUs because its expectation-maximization algorithm is poorly suited for dereplicated data. Finally, ACT applies a decision tree to reconcile the Emu, Sintax, and LACA outputs (Fig. S1), assigning the lowest well-supported rank as the final consensus taxonomy for each read. Importantly, if the confidence of the Sintax assignment is below a user-adjustable threshold, designated “NA” for “not assigned”, taxonomy is not assigned for that rank to avoid overclassification. The LACA OTU number is included within the taxonomic assignment to enable diversity tracking at any taxonomic level (e.g. *Bacillaceae sp*. OTU 1 or *Bacillus sp*.OTU 2).

Following TaxId assignment, either directly from the database or generated using the TaxonKit create-dump tool (16) for databases lacking TaxIds, ACT tallies assigned TaxIds by sample and outputs a final consensus abundance table containing the read counts for each TaxId with corresponding OTU. For users focused only on named taxa, ACT also outputs a read count table of TaxIds that excludes the OTUs. The tables are unfiltered to avoid omitting rare taxa, leaving application of minimum abundance thresholds to user discretion.

### Formatting standard databases

We used the following databases: Greengenes2 (version 2024.09); Genome Taxonomy Database (GTDB) (release 226); GSR-DB (GSR-DB_full-16S accessed 6/11/2025), which integrates Greengenes, SILVA, and RDP sequences (17); Emu-DB (2020 build), which combines the bacterial and archaeal 16S rRNA sequences from the NCBI 16S rRNA RefSeq database and the prokaryotic Ribosomal RNA Operon Copy Number Database (rrnDB)(18); and an updated version of the Emu-DB (Emu25-DB) that we generated in 2025 (NCBI and rrnDB accessed 11/1/2024). Database creation and formatting is further described in the Supplemental Methods. Characteristics and metrics of each database are detailed in Table S1.

### ACT database (ACT-DB) construction

The ACT-DB is a sequence-similarity-aware version of Emu25-DB, integrating RefSeq and rrnDB sequences. It was built using the grouping function of the ACT database builder (database_builder.sh --group), which globally aligned all Emu25-DB sequences using Minimap2 to identify those sharing >99.5% identity. This threshold represents the midpoint between PHRED Q-scores of 20 and 30, corresponding to the upper accuracy range currently observed in ONT sequencing. Above 99.5% identity, differences may reflect sequencing error rather than biological variation. Following exclusion of outliers from suspected contamination, sequences sharing >99.5% identity were clustered into named groups (e.g., *Bacillus* species group 1). The grouping script also outputs a list of species within each group and a list of sequences assigned to each group.

### Mock community standard

The ZymoBIOMICS Microbial Community DNA Standard D6305 (Zymo Research) (hereafter “ZymoCom”) contains genomic DNA of eight bacterial species and two fungi, with theoretical species abundances and 16S rRNA sequences for each member specified by the manufacturer. This study used ZymoCom from two different lots (ZRC195708 [ZymoCom A-D] and 29120 [ZymoCom E-H]), with each sequenced as part of 4 multiplexed libraries.

### Generation of simulated ZymoCom reads

Based on the eight bacterial ZymoCom members, we created three synthetic data sets, each comprising five samples: (i) equal species abundances, (ii) equal species abundances plus synthetic spike-in Ec5001 reads, and (iii) unequal species abundances varying 10^5^-fold between lowest and highest abundance reads. Genomes for the eight species were downloaded from Zymo Research documentation and *in silico* 16S rRNA gene PCR performed with AmpliconHunter v2.0 (19). Simulated ONT 16S rRNA output was generated for each genome using Seq2Squiggle (20) v0.3.4 (predict -n 100000 -r 1600 --duration-sampler False --noise-std 2.0) and basecalled using Dorado (dna_r10.4.1_e8.2_400bps_sup@v5.0.0). Seqkit (21) v2.10.0 was used to randomly select simulated reads (shuffle, sample -p, head) of each taxon and combine them to create a sample. The number of reads of each taxon in a sample was dependent on dataset and R was used to add between and within sample variation to the initial counts. Relative abundances for each sample were multiplied by a random number to simulate variation in sequencing depth between samples and then the command “jitter” was used to add random noise to each abundance within a sample. Detailed scripts are provided on GitHub. Table S2 shows the number of simulated reads for each species in each sample.

### MAize Rhizosphere Synthetic community (MARSc)

MARSc is comprised of the model rhizosphere colonist *Pseudomonas putida* KT2440 and 30 bacterial strains isolated from the maize root environment (22–27). Thirteen of the 30 isolates are novel species that were absent from the reference databases when the analyses were performed, but have since been deposited in RefSeq (27). For select analyses, the 16S rRNA genes for the 30 isolates were manually added to the ACT-DB, with unique, arbitrary TaxIds assigned for novel species and existing TaxIds used for higher ranks of novel species and all ranks of known species.

### MARS inoculation of maize and rhizosphere microbiome sample collection

Surface-sterilized and pre-germinated maize seeds were inoculated with a dilute suspension of the MARSc community prior to planting in cone-tainers filled with a 75:25 soil:sand mix. Plants were fertilized at planting, grown in a growth chamber, and rhizosphere samples collected 21 days after planting. Rhizosphere soil was collected from the root by washing with sterile phosphate buffer and then the washates were centrifuged and the pellets stored at −80°C. DNA was extracted from the pellets using a Qiagen PowerSoil Pro 96-well plate DNA extraction kit, with plates loaded using a modified method to decrease cross-contamination (28). A more detailed description of these steps is available in the Supplemental Methods.

### Generation and sequencing of barcoded full-length 16S rRNA amplicon libraries

Amplicon libraries were generated using custom primers (Integrated DNA Technologies) containing 5’ 13-bp barcodes (29) fused to degenerate 27F (GAGTTTGATYMTGGCTCAG) and 1492R (CGGYTACCTTGTTACGACTT) 16S rRNA primers. Library and barcode details are provided in Table S3. PCR reactions (50 µL) contained the following: 1X KAPA HiFi Fidelity buffer, 300 μM of each deoxynucleotide triphosphate, 0.5 units KAPA HiFi HotStart Polymerase (all from Roche), 5% v/v DMSO, spike-in mix, 100 nM each of barcoded 27F and 1492R primers, and 5 ng community DNA (ZymoCom or rhizosphere DNA). Spike-in mix details are provided in the Supplemental Methods. PCR cycling conditions were as follows: 95°C for 5 minutes, 25 cycles of 45 sec at 98°C, 45 sec at 54°C, and 100 sec at 72°C, and a final extension of 72°C for 5 min. Additional information on amplicon library generation is available in the Supplemental Methods. PCR products were pooled into libraries at roughly equal concentrations, as estimated from gel band intensities, and purified using a Monarch PCR & DNA cleanup kit (NEB). ONT sequencing was performed at SeqCenter (Pittsburg, PA), with library preparation using the PCR-free ONT Ligation Sequencing Kit (SQK-NBD114.24) with NEBNext® Companion Module (E7180L), sequencing using R10.4.1 flow cells in 400-bps sequencing mode with minimum read length of 200 bp and no adaptive sampling, and super-accurate base calling using Guppy v6.5.7 (model dna_r10.4.1_e8.2_400bps_modbases_5mc_cg_sup.cfg) with no quality trimming.

### Read processing

Raw reads were inspected for quality and size distribution using NanoPlot v1.44.1 (14). Based on these distributions, ACT pre-processing was performed with chopper v0.10.0 using a minimum read length of 800 bp, maximum read length of 4,000 bp to preserve concatenated reads, and a minimum average Q-score of 12. The 13-bp barcodes used in this study each differed by at least two bases, so demultiplexing and barcode trimming with Cutadapt v5.1 was performed using a minimum barcode overlap of 12 and a maximum error of 1 base. Demultiplexed reads were then inspected again and filtered using a minimum length of 1300 bp, maximum length of 1700 bp, and minimum average Q-score of 15 based on the observed quality and size distribution and expected amplicon size. Filtered read statistics, as determined using Seqkit (21) v2.10.0 (stats -aT), are provided in Table S3. Filtered reads were classified with ACT, Emu v3.5.1, and Sintax v2.30.0 using indicated reference databases. For ACT, the NA threshold, which corresponds to sintax bootstrap confidence values, was set at 0.1 for ZymoCom samples and 0.3 for rhizosphere samples. Emu and Sintax outputs were processed in R to generate read count tables (detailed scripts in supplemental methods); ACT read count tables were produced via pipeline scripts.

### Downstream data analysis

Abundances were analyzed using the R package microeco (30). No additional abundance thresholding or rarefaction was applied. The microeco taxonomy table was revised with descriptive placeholders for unassigned ranks by appending the lowest identified rank followed by “unknownRank”, (e.g., Bacillaceae_unknownGenus_sp.). The percentage of classified reads was calculated as the number of classified reads per sample-pipeline-database combination divided by that sample’s total reads. For species richness, reads that were classified to the species level had their OTU, which would correspond to sub-species, removed so that all classifications were at the rank species or higher. Chao1 and Observed species richness indices were calculated using microeco, with each index and database pair calculated separately, and, within each index-database pair, the pipelines compared by Kruskal-Wallis test followed by Dunn’s post-hoc test. The difference between expected and observed read counts in the simulated mock community samples were calculated for each taxon as percent error with the following equation 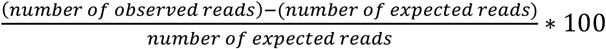. Precision, recall, and model accuracy (F1 score) were calculated using presence/absence data at each level of taxonomy. Precision of positive identifications was calculated as 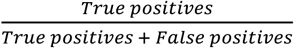. Recall, which reflects the potential for identifying false negatives, was calculated as 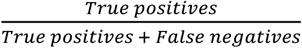. Model accuracy (F1 score) is the harmonic mean of Precision and Recall and is derived as: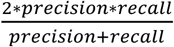. A higher F1 score reflects better model accuracy.

### Access and usage

The ACT pipeline, the ACT-DB, and documentation are publicly available via GitHub at https://github.com/Halverson-lab/Amplicon_Consensus_Taxonomy.

## RESULTS

### ACT performs comparably or better than Emu or Sintax for profiling a simple mock community

We first assessed ACT pipeline performance by comparing the theoretical versus observed composition of the eight-member “ZymoCom” mock community when profiled with 16 pipeline– database combinations. These included three pipelines (ACT, Emu, and Sintax) paired with four databases (Emu25-DB, GSR-DB, Greengenes2, and GTDB), plus Emu with its original 2020 database (Emu-DB), which was included as a benchmark. Overall, ACT, Emu, and Sintax classified more reads at the genus than the species level, with ACT and Emu outperforming Sintax in both taxon detection accuracy (i.e., identifying known taxa) and abundance estimation (Fig. 2).

**Figure 2.**
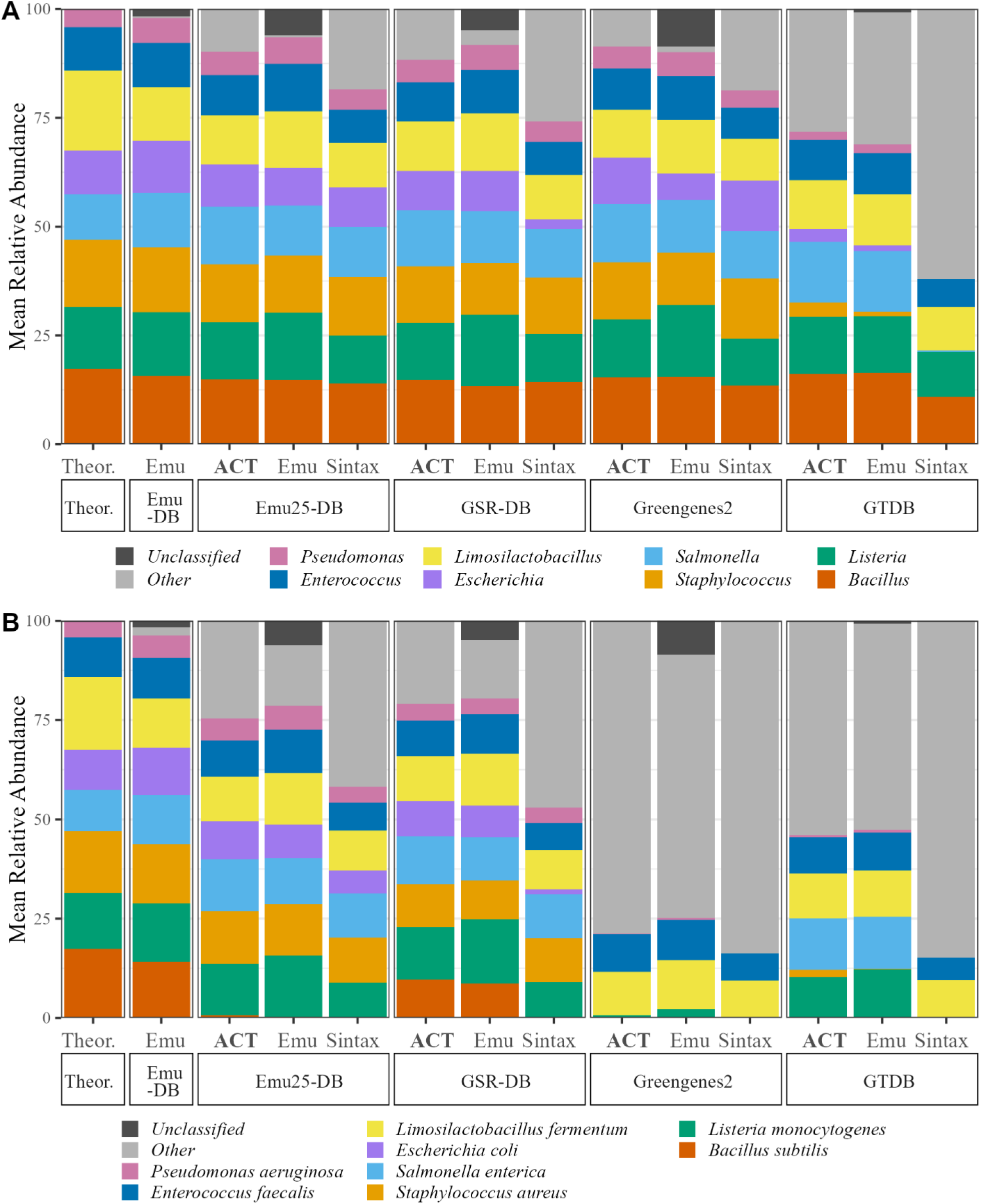
Factorial assessment of pipeline and database effects on relative abundance profiles of the Zymo mock community at genus (A) and species (B) levels. “Other” includes all reads that were not identified as a specific community standard at the genus (A) or species (B) level. The x axis identifies the database (DB) in the boxes below pipelines used. Theor. denotes the theoretical composition of the Zymo mock community according to Zymo Research documentation. The y axis is the mean relative abundance (n = 8).

Detection accuracy varied substantially across reference databases. With GTDB, multiple taxa were underrepresented or completely undetected, even at the genus level. Closer inspection indicated that GTDB classification failures stemmed, at least in part, from contaminated assemblies, as illustrated by the misclassification of *E. coli* reads due to the presence of *E. coli* sequences in the metagenome assembly of *Alloprevotella* sp004555055. Greengenes2 produced genus-level profiles similar to expected, particularly when paired with the ACT and Emu pipelines, but missed multiple taxa at the species level. Detection failures were due to both nomenclature differences, as illustrated by annotation of *Salmonella enterica* as *S. bongori*, and database gaps, as shown by the absence of annotated *E. coli* sequences. Because of these low detection accuracies, GTDB and Greengenes2 were excluded from subsequent analyses.

When paired with Emu25-DB or GSR-DB, both the ACT and Emu pipelines classified >88.5% of reads as expected genera at near theoretical abundances. This was lower than the 97.8% of reads classified by Emu when paired with Emu-DB, which is a considerably smaller database. Of note, reads that were not classified as expected genera were handled differently by the two pipelines: ACT classified most “other” reads to higher taxonomic ranks while Emu effectively discarded them by leaving them unclassified (Table S4). Thus, ACT preserved more taxonomic information across the dataset. When using ACT with the Emu25-DB, the abundance of reads for the *Bacillus* genus was at near-expected levels, but few *Bacillus* reads were assigned at the species level. In contrast, Emu classified *Bacillus* reads at the species level, but these assignments varied based on the database used. For example, the reads classified by Emu as *B. subtilis* with Emu-DB or GSR-DB were classified as *B. spizizenii* with Emu25-DB. Although the difference between *B. subtilis* and *B. spizizenii* reflects nomenclature updates, nomenclature changes did not explain why most *Bacillus* reads were not assigned to a species by ACT with the Emu25-DB. This prompted us to further investigate *Bacillus subtilis* 16S rRNA gene sequences.

*B. subtilis* has ten 16S rRNA genes, and BLAST searches revealed that these ten genes share >99% identity with 16S sequences from five other *Bacillus* species, including *B. spizizenii*, with most sequences differing by <5 bp. Even with high-accuracy ONT reads, this minimal variation is insufficient to reliably distinguish among these *Bacillus* species. Thus, although Emu appeared to outperform ACT by assigning *Bacillus* reads to a species, this reflects overclassification rather than improved accuracy. In contrast, by not assigning *Bacillus subtilis* reads to a species, ACT with Emu25-DB captured the inherent ambiguity in *Bacillus* 16S rRNA gene sequences.

### The ACT-DB accounts for sequence ambiguity to improve classification accuracy

Because long-read amplicons are intended to improve resolution at lower taxonomic ranks, we sought an approach to attain as much resolution as possible even when full-length 16S rRNA sequences could not distinguish closely related species. This led us to build the ACT-DB, a database in which 16S rRNA reference sequences with >99.5% similarity are clustered into multi-taxa groups that are assigned as a group rather than as individual species (e.g., *Bacillus-Calidifontibacillus spp*. group 299 contains *B. subtilis* and *B. spizizenii*). Of the 4371 genera represented in Emu25-DB, 1406 had 16S rRNA sequences that clustered into multi-taxa groups, including all ZymoCom genera except for *Limosilactobacillus fermentum*. Of the 1406 genera with sequences in multi-taxa groups, 1102 also had 16S rRNA sequences that did not cluster into groups. Thus, the resolution afforded by 16S rRNA sequences can vary across gene copies within the same species, and the unique 16S rRNA copies in many of these species can enable unambiguous species detection.

Classification of the ZymoCom samples using ACT-DB led to greater agreement between the three pipelines at the species level. For example, whereas with Emu25-DB Sintax failed to classify any *Bacillus* reads to the species level and Emu identified more than 3 *Bacillus* species, with ACT-DB all three pipelines detected a similar abundance of reads belonging to the *Bacillus-Calidifontibacillus spp*. group 299 (Fig. 3). ACT-DB also preserved species level classification where possible, as illustrated by the successful classification of *L. fermentum* (Fig. 3). By accounting for sequence ambiguity, the ACT-DB limits spurious resolution in favor of accurate, reliable classifications.

**Figure 3.**
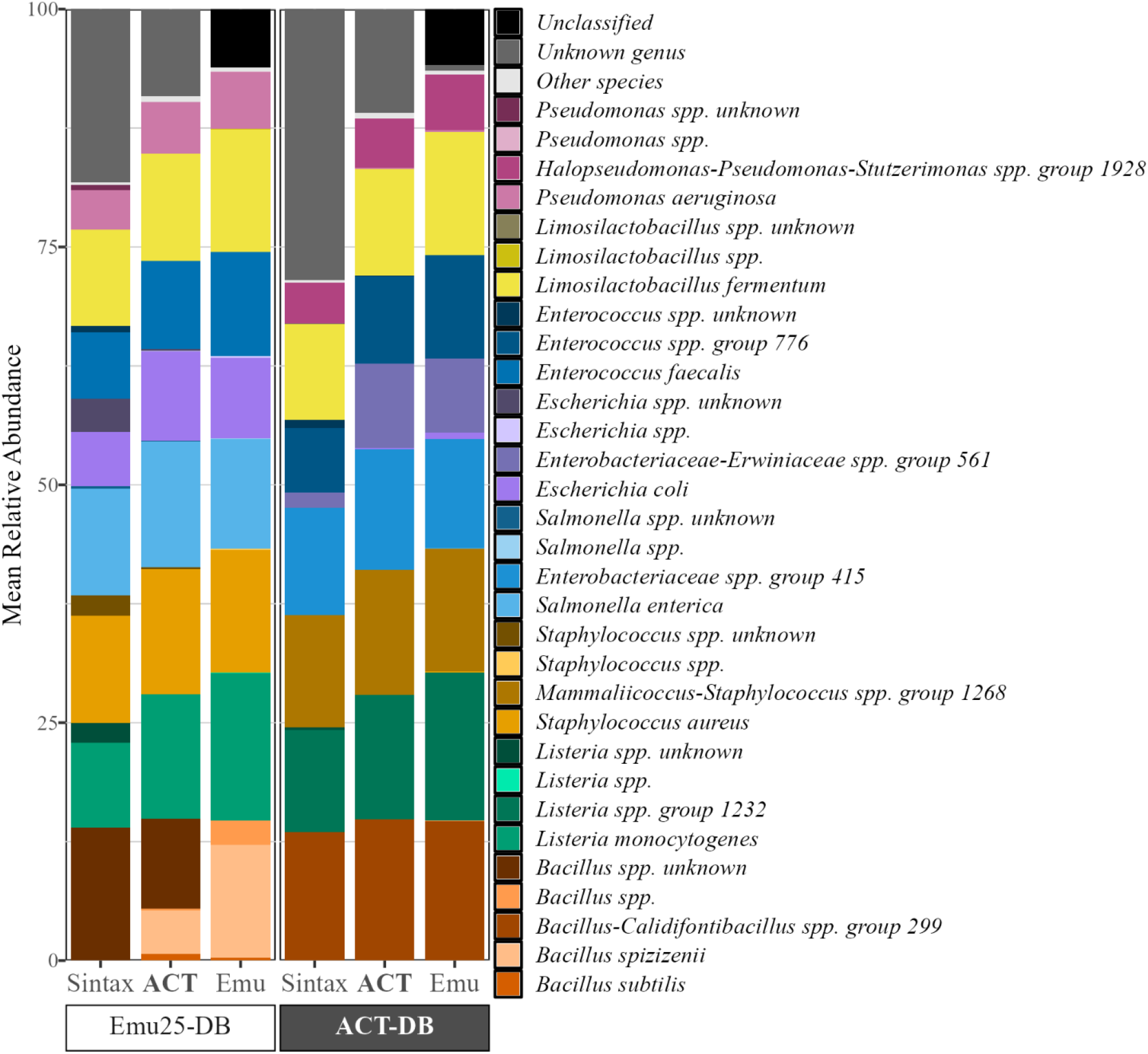
Effect of sequence-based clustering on the taxonomic classification of the Zymo mock community using the ACT, Emu, and Syntax pipelines. The x axis identifies database (DB) in the boxes below pipelines used. The y axis is the mean relative abundance (n = 8). All reads assigned to a species not belonging to the ZymoCom are grouped under “Other species”. Unclassified designates reads that were not assigned a genus or species classification. Unknown genus designates reads that are classified at the family level or higher. Species listed as ‘Genus sp unknown’ were classified only to the genus level.

### ACT outperforms Emu in profiling synthetic communities with rare and novel taxa

We also benchmarked ACT performance using a synthetic dataset comprised of 16S rRNA gene sequences extracted from the genomes of the bacterial ZymoCom members. We incorporated the synthetic DNA spike-in Ec5001 as a standard for improving 16S rRNA amplicon profiling (30). To determine a pipeline’s ability to handle species not present in the database, we first compared the results from simulated samples comprised of equal taxon abundance with the ACT and Emu pipelines using ACT-DB with and without Ec5001 included. ACT and Emu pipelines performed similarly to the theoretical relative abundances when Ec5001 was present in ACT-DB, but Emu’s accuracy diminished greatly when Ec5001 was omitted from the database, generating inflated abundances for 6 of the 8 genera (Fig 4). Closer examination of Emu results using ACT-DB without Ec5001 shows that Emu classifies near the same number of reads for each genus but discards unclassified Ec5001 reads entirely (Fig 4); the discarded reads are not included in the relative abundance calculations which artificially inflates the relative abundance of the remaining species. In contrast, the relative abundances estimated by ACT closely resembled theoretical values regardless of if Ec5001 was included in the database or not and identified most Ec5001 reads to a single OTU. These results demonstrate the strength of ACT, in concert with the ACT-DB, in maintaining accurate composition estimates for communities containing unknown taxa (Fig. 4).

**Figure 4.**
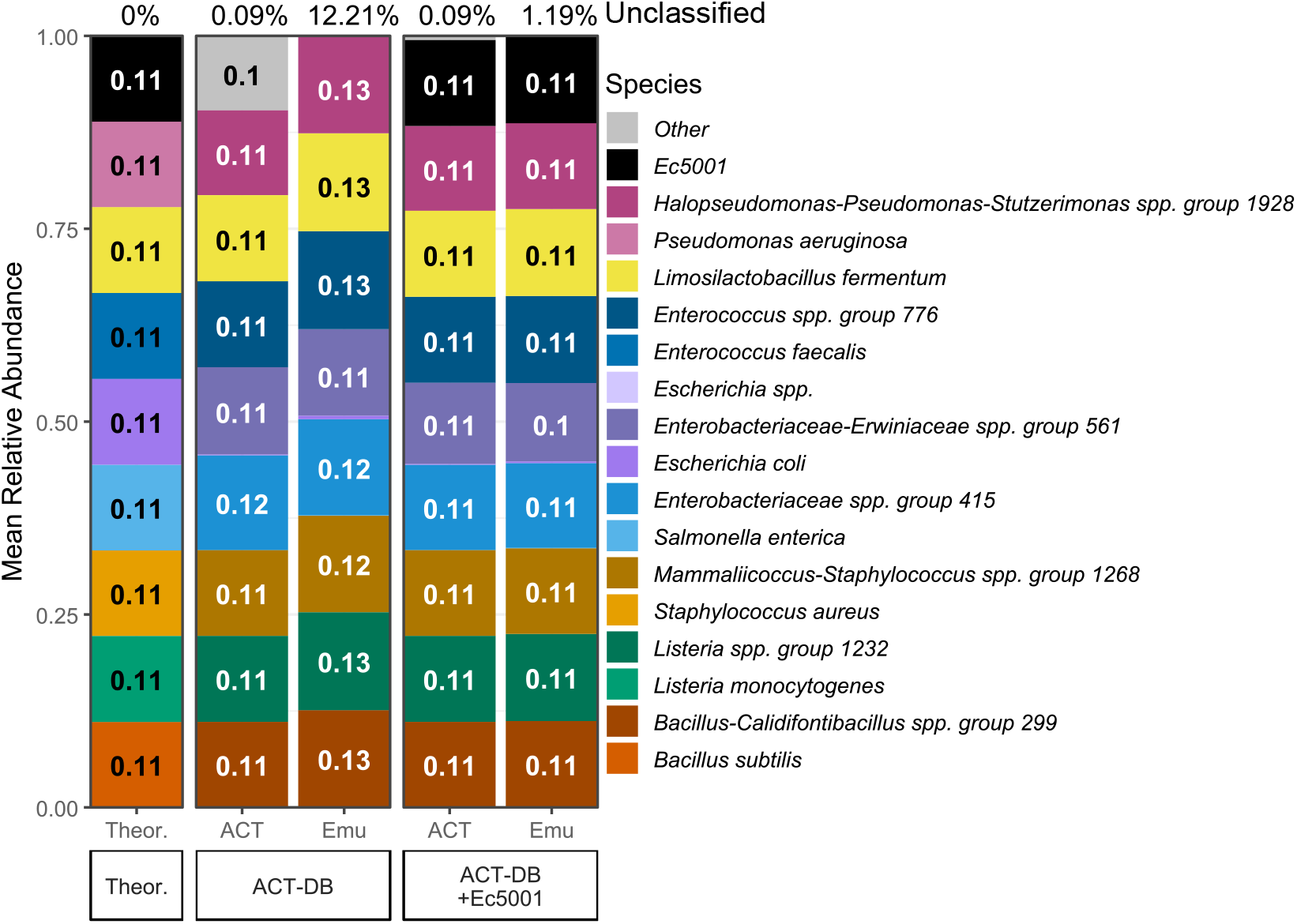
Detection of novel taxa in a simulated mock community by ACT or Emu. Theoretical is the expected relative abundance based on equal representation of each taxon in the simulated read counts. The x axis identifies which pipeline (ACT or Emu) was used and whether synthetic spike-in Ec5001 was included in the ACT-DB. The y axis is the mean relative abundance (n = 5). Listed above each bar is the mean (n=5) percent of reads that were left unclassified and hence were excluded from the relative abundance calculations.

To evaluate detection of rare or low-abundance taxa, we compared ACT and Emu performance using two simulated datasets comprising samples with either equal (~ 11% each) or unequal (~0.00046% to ~46%) abundances. For each analysis, we determined the percent error between the observed vs expected read counts for each of nine taxa. ACT outperformed Emu across both datasets. With the equal abundance dataset, ACT achieved equal or lower error for all taxa (Fig 5). With the unequal abundance dataset, ACT yielded dramatically lower error than Emu for three taxa (Fig 5), including the two least-abundant taxa that Emu failed to detect entirely, likely due to Emu’s built-in abundance thresholding. Although both ACT and Emu detected Ec5001, the detection error was only 0.42% for ACT compared to 80% for Emu (Fig 5). ACT and Emu yielded more comparable percent error for the other six taxa. Overall, these data demonstrate that ACT better captures rare taxa, preserving more information about community composition and structure than Emu.

**Figure 5.**
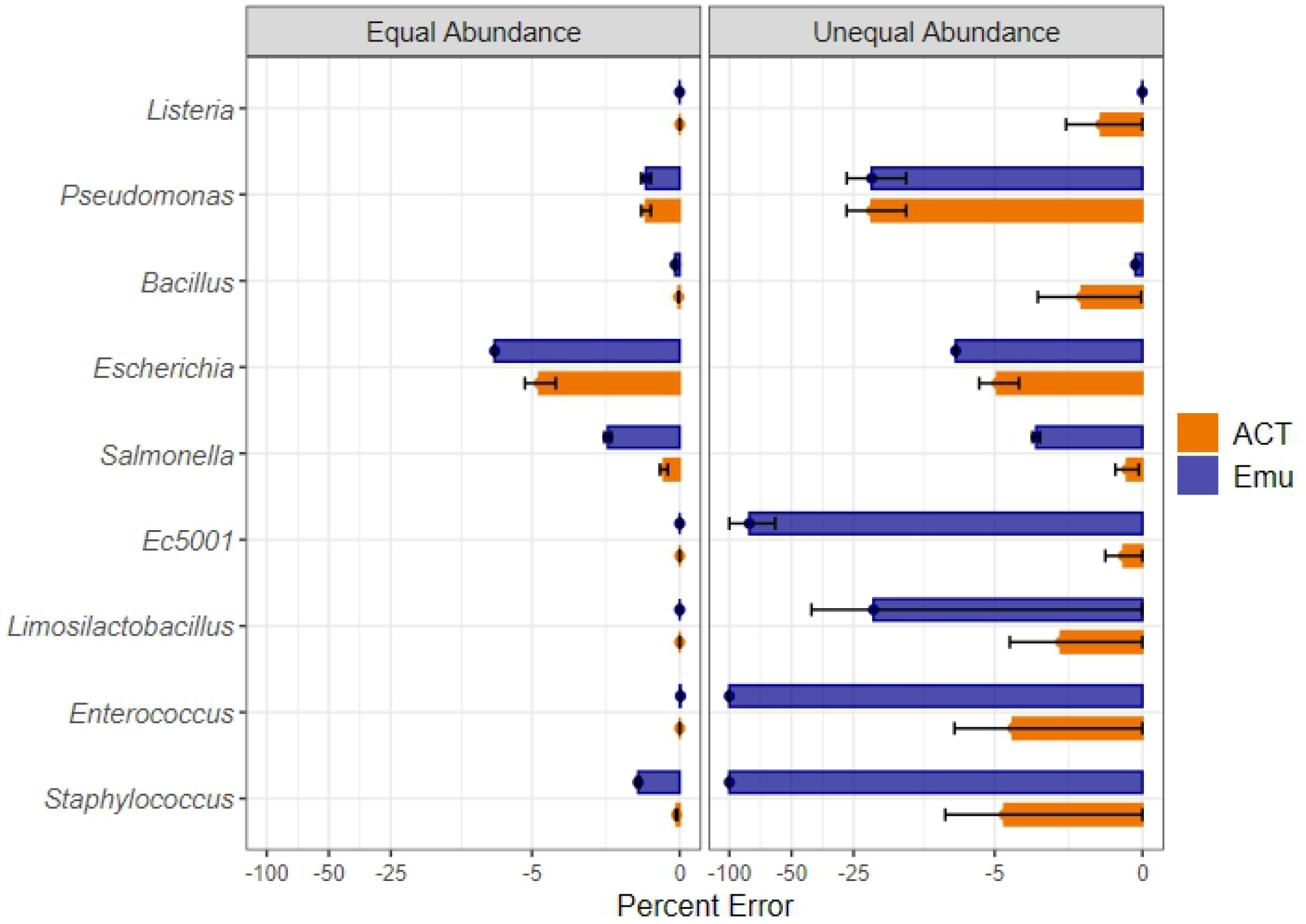
Influence of pipeline on genus-level composition error of a simulated mock community. Values represent the mean (n=5) ± standard error of the percent error of the mock community with the synthetic spike-in standard Ec5001 included in the ACT-DB. Taxa are listed in order of abundance in the unequal abundance data: *Listeria* (~46%), *Pseudomonas* (~46%), *Bacillus* (~4.6%), *Escherichia* (~2.3%), *Salmonella* (~0.46%), Ec5001 (~0.046%), *Limosilactobacillus* (~0.046%), *Enterococcus* (~0.0046%), *Staphylococcus* (~0.00046%).

We also used the simulated datasets to examine ACT and Emu performance in terms of precision (reliability of positive identifications), recall (sensitivity), and overall model accuracy (F1 score) (Fig 6). Regardless of pipeline, precision and model accuracy were better with ACT-DB than with Emu25-DB, particularly at the genus and species ranks. Although recall at the species rank was lower with ACT-DB than Emu25-DB, this was expected because most ZymoCom members have ambiguous 16S sequences that cluster into multi-species groups within the ACT-DB. Thus, lower recall at the species level reflects a deliberate reduction in overclassification in favor of increased precision and accuracy.

**Figure 6.**
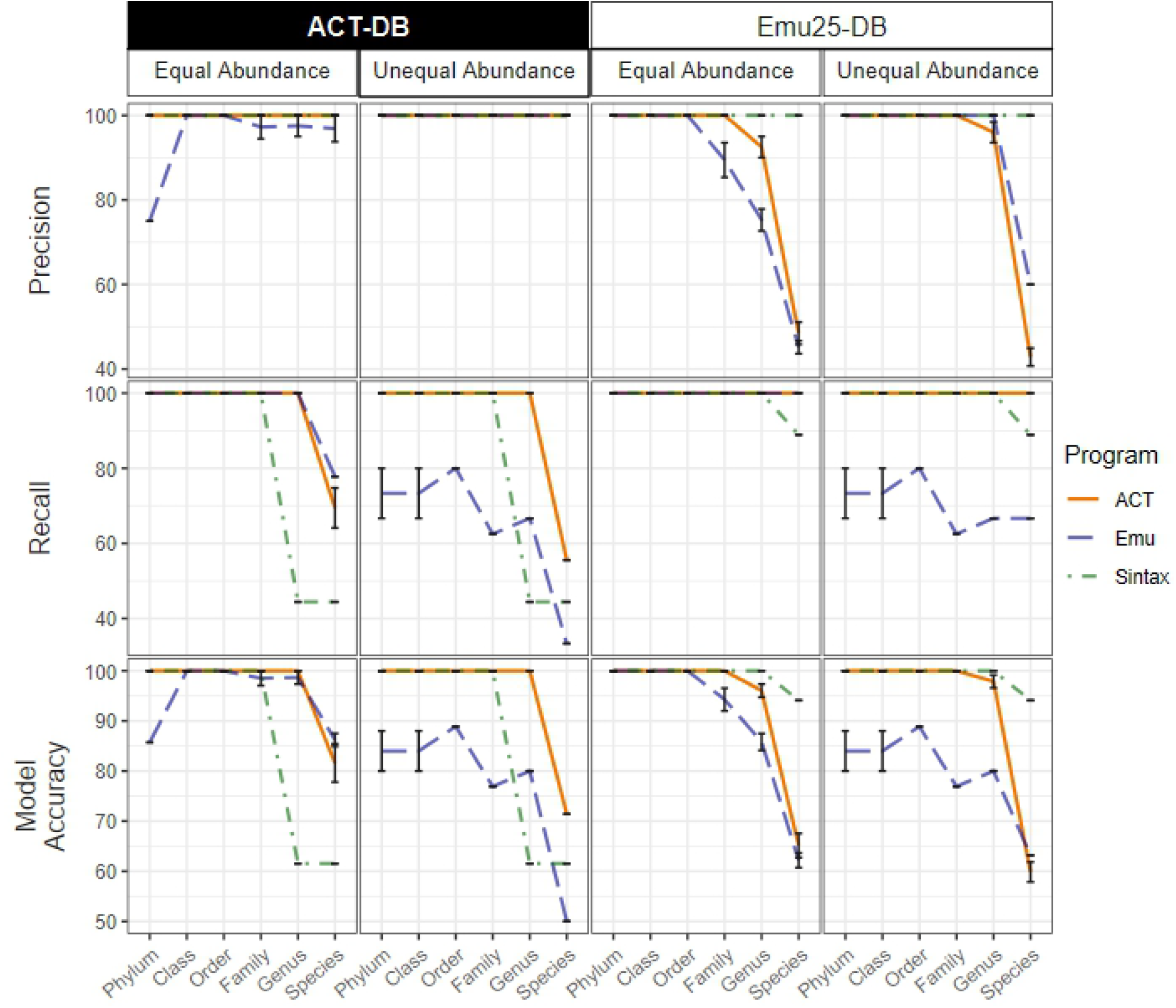
Precision, recall, and model accuracy (F1 score) of ACT and Emu pipelines using ACT-DB or Emu25-DB when profiling simulated datasets comprising samples with equal or unequal abundances of nine taxa. Lines show the mean (n=5) and error bars denote the standard error of the mean.

### ACT captures more diversity in complex natural communities than Emu or Sintax

We also compared pipeline performance using a complex maize rhizosphere community that presumably contains novel species. Emu classified an average of 87% of reads per sample, but as little as 47% in some samples, reflecting its all-or-nothing approach of assigning reads to the nearest species or not at all (20). In contrast, both ACT and Sintax classified nearly 100% of reads, but ACT classified an average of 71.8% of reads to the genus rank and Sintax only 33.6%. An average of 65.1% of reads were assigned OTU classifiers by the ACT pipeline with the ACT-DB, indicating that ACT captured likely novel species absent from the database. Additionally, ACT reported significantly higher species richness metrics than Emu or Sintax (Fig. 7). While the species richness appears particularly high, our sequencing depth was sufficient to support it, with an average of 189,037 reads per sample (Table S3Thus, ACT reported greater richness and diversity in rhizosphere samples, primarily by capturing rare and novel species.

**Figure 7.**
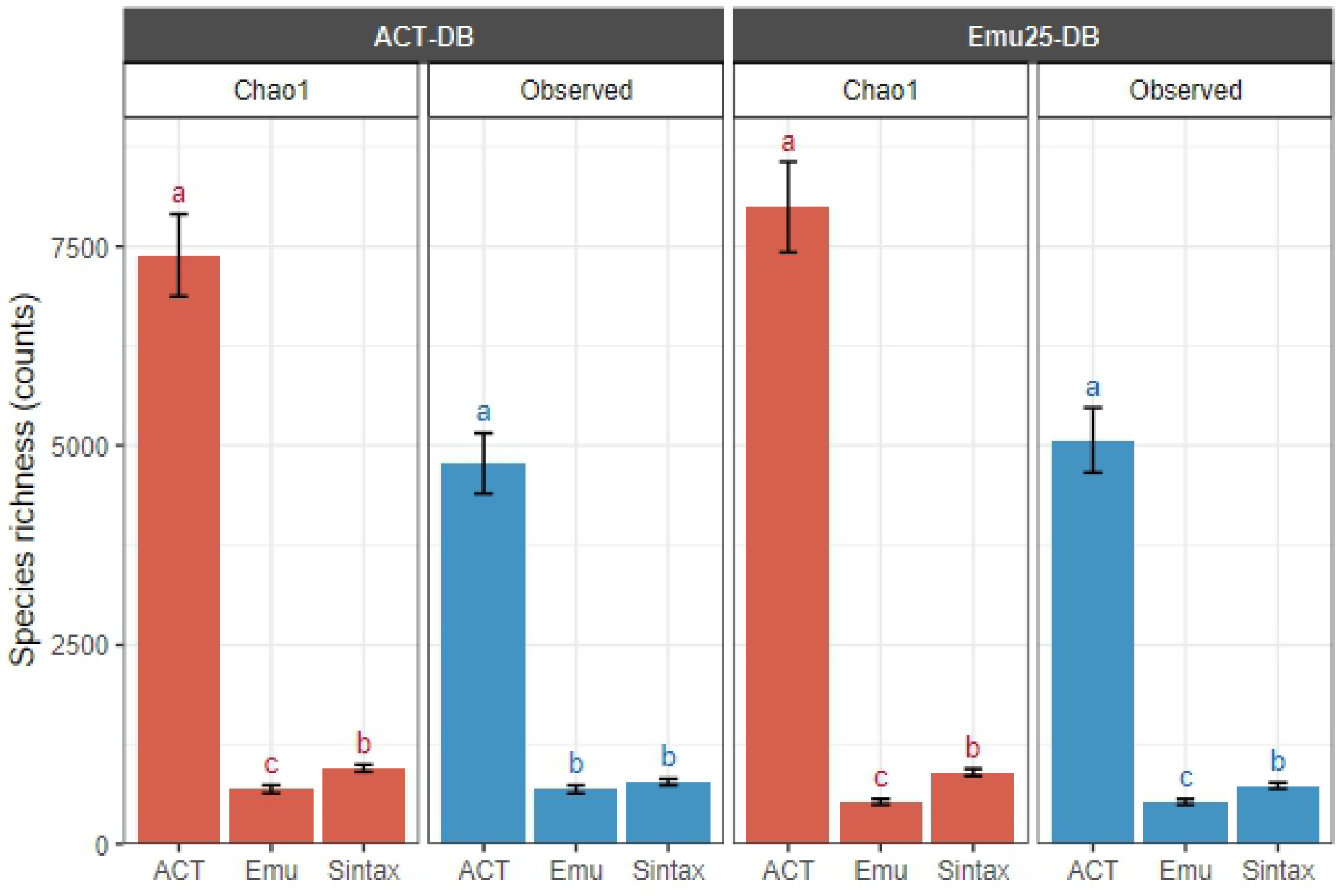
Species richness of natural maize rhizosphere communities is affected by pipeline and reference database. Maize seeds were inoculated with the MARSc community and grown in field soil. For reads that were not assigned to a species the lowest taxonomic rank and OTU were used instead. Values represent the mean (n=5) ± standard error of the mean. A separate Kruskal-Wallis test with Dunn’s post-hoc test was performed for Chao1 and Observed species richness for each database. Letters above a bar denote significant (p < 0.05) differences between pipelines for that metric and database.

### ACT-DB decreases misclassification of novel taxa in complex natural communities

The maize rhizosphere community data also provided insights into how ACT accommodates the detection of novel species. At planting, maize was inoculated with the MARSc synthetic community, which contains 13 potentially novel species that were not represented in the reference database at the time of this study. For illustration purposes here, we selected *Sphingobium* R-21 to represent a novel species due to its abundance and the fact that relatively few other *Sphingobium* species were present. We observed a high degree of similarity in the 16S rRNA sequences amongst *Sphingobium* species, with R-21 classified as a member of *Sphingobium spp. group 2200*. As a consequence, ACT with Emu25-DB classified a majority of reads as *S. fuliginis* (Fig. 8). Adding R-21 sequences to the database improved detection, but the similarity amongst species led to a decrease in sensitivity, observed as an increase in the proportion of reads classified as *S. sp. unknown*. Use of ACT-DB, both with and without R-21 sequences added, led to more consistent results, with both reporting near equal abundances of *Sphingobium spp. group 2200* (Fig. 8). This consistency reflects the power of ACT-DB to more accurately classify species that belong to groups with low inter-species variation.

**Figure 8.**
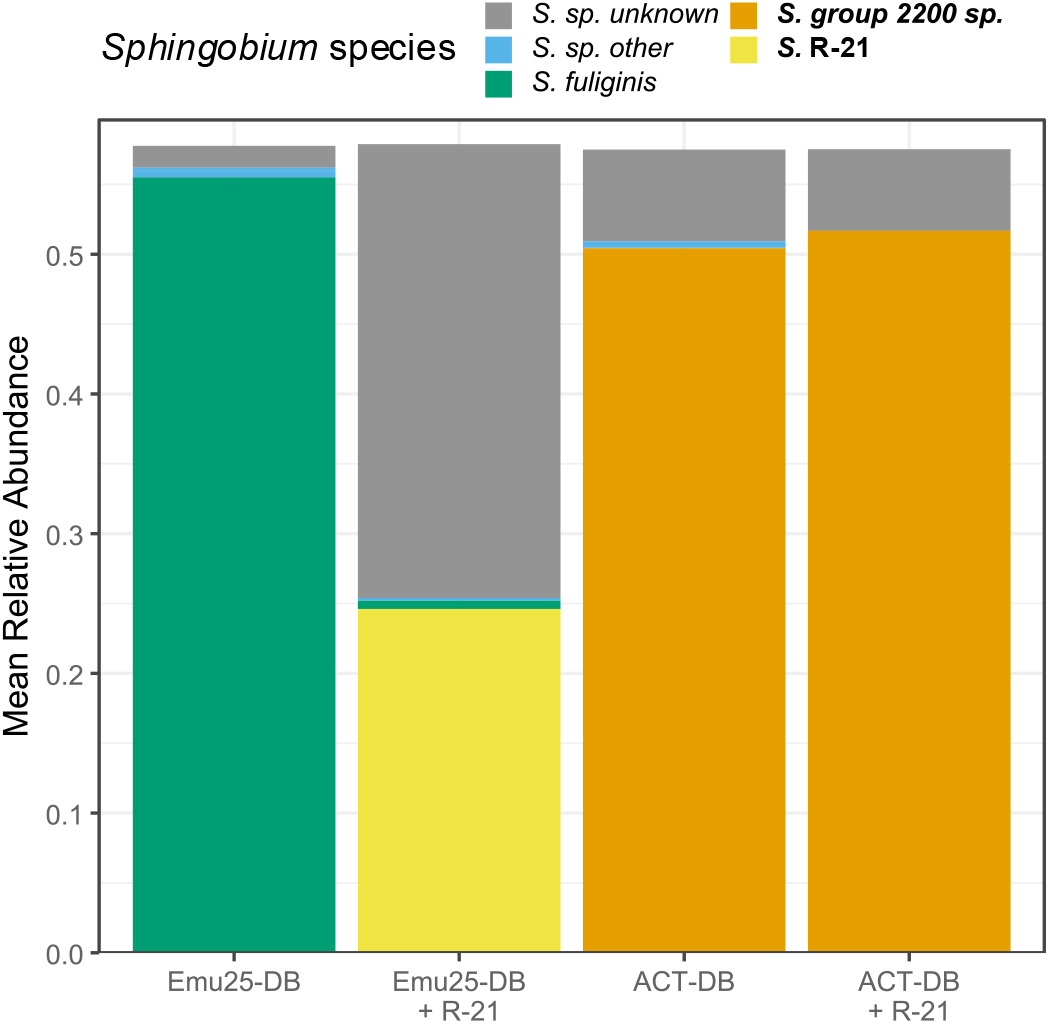
Effects of sequence-based clustering on detection of *Sphingobium* R-21 in a maize rhizosphere community using the ACT pipeline. Mean (n=5) relative abundance of all *Sphingobium* species detected in the rhizosphere of maize inoculated with MARSc (including *Sphingobium* R-21) and grown in field soil. Species “unknown” denotes reads classified to the genera *Sphingobium* and not classified to a species. Species “other” combines all *Sphingobium* species except *S. fuliginis, S*. R-21, and *S. spp. group 2200*. “+R-21” means all annotated 16S rRNA sequences from *S. R-21* were added to the reference database. ACT-DB was created by removing the *S*. R-21 sequences from ACT-DB+R-21.

## DISCUSSION

As ONT sequencing becomes more accurate and affordable, robust workflows are needed to translate long-read amplicons into community profiles with high resolution and accuracy. The ACT pipeline provides a robust workflow by integrating outputs from Emu, Sintax, and LACA to generate consensus taxonomic assignments for individual reads. Through its decision tree, ACT leverages the complementary strengths of each individual tool: (i) the ability of Emu to classify reads from known species with high accuracy; (ii) the ability of Sintax to minimize overclassification, including reads from rare and unknown taxa, and provide bootstrap-associated assignments at all ranks; and (iii) the ability of LACA to capture intra-taxon (e.g., intra-genus) diversity for unknown species by clustering sequences into OTUs. By reconciling taxonomic information across these tools, ACT reduces misclassification and retains all valid reads in a dataset, improving diversity estimates, which is particularly important in complex communities.

Although the consensus approach of ACT provides clear benefits over single-classifier approaches, ACT performance is strongly influenced by the reference database. A database with misnamed sequences or outdated taxonomy leads to classification errors. Databases with broad taxonomic coverage will support classification of a greater number of reads if represented in the database. However, less intuitively, a database with broader coverage can also increase ambiguity in classification when sequences are not sufficiently different to resolve closely related taxa.

Consequently, smaller, less comprehensive databases can both reduce species-level resolution by omitting taxa and artificially inflate it by overlooking ambiguous sequences. This effect was evident in our ZymoCom profiles, where *Bacillus* reads either were classified as distinct species or were not speciated at all, depending on the reference database. Sequence ambiguity that leads to misclassification distorts the resulting community profiles.

We are not the first to recognize that full-length 16S rRNA gene sequences do not always afford species-level resolution (31), but we propose a new approach to address this issue: a reference database (ACT-DB) that clusters highly similar reference sequences into multi-taxa groups. By explicitly accounting for sequence ambiguity, ACT-DB improves precision, and by defining the members in each group, ACT-DB preserves taxonomic information that can facilitate interpretation and downstream analyses. We found that community profiles were more consistent across pipelines when using the ACT-DB compared to the non-grouped Emu25-DB, suggesting that inconsistent classification of ambiguous sequences may contribute to variation between studies.

Collectively, our results indicate that the ACT pipeline and ACT-DB are powerful tools for precise, high-resolution profiling of microbial communities using long-read amplicons. We designed ACT to be both accessible and flexible, integrating automated execution and user-defined settings for read processing and classification. Also, the ACT decision tree provides a framework that can be modified or extended with new tools in future iterations. The ACT-DB is equally adaptable, allowing users to readily add sequences, update the entire database, or substitute custom databases as needed. Additionally, by using NCBI TaxIds and accessions, ACT and the ACT-DB ensure that previous classifications can be easily updated to reflect taxonomic changes. Finally, though tested here for performance using 16S rRNA amplicons, ACT can be used with any marker region when provided a suitable reference database, be it fungal ITS regions, eukaryotic 18S rRNA genes, or longer multi-gene regions (11). We encourage users to share feedback through the ACT GitHub repository to guide future improvements and ensure the continued utility and broad applicability of ACT and the ACT-DB.

## DATA AVAILABILITY

The ZymoCom and rhizosphere sequences have been deposited in the Sequence Read Archive under accession no. PRJNA1442200; individual accession numbers are listed in Table S3. The simulated reads, reference databases, and code for analyses are available on Open Science Framework (https://doi.org/10.17605/OSF.IO/TJCW4). The ACT pipeline is available on GitHub (https://github.com/Halverson-lab/Amplicon_Consensus_Taxonomy).

## ACKNOWLEDGMENTS

This research was supported by a USDA National Institute of Food and Agriculture AFRI (grants 2019-51300-30248, 2021-67019-34833, and Hatch project IOW04108) and the National Science Foundation (grants DRL-1814001 and DGE-1545453).

