## Supplemental Information for "A Bioinformatic Pipeline for Consensus Taxonomic Classification of Long-Read Amplicons"

### SUPPLEMENTAL METHODS

#### Formatting standard databases

This study used the following standard databases: Greengenes2 (version 2024.09); Genome Taxonomy Database (GTDB) (release 226); GSR-DB (GSR-DB_full-16S accessed 6/11/2025), which integrates Greengenes, SILVA, and RDP sequences (1); Emu-DB (2020 build), which combines the bacterial and archaeal 16S rRNA sequences from the NCBI 16S rRNA RefSeq database and the prokaryotic Ribosomal RNA Operon Copy Number Database (rrnDB)(2); and Emu25-DB (NCBI and rrnDB accessed 11/1/2024), which is an updated version of the Emu-DB built in 2025 and generated using the steps outlines in the Emu documentation. Greengenes2, GTDB, and GSR-DB were converted to Emu-formatted databases using Emu's "build-database" command and subsequently to Sintax-formatted databases using emu_to_sintax_db_converter.R script. Scripts for generating all databases are provided in the ACT documentation.  For database performance comparisons, taxonomy files for all databases were merged to create a master table of all TaxIds and corresponding lineages, with formatting to correct and simplify naming conventions, including updating phyla names, removing special characters and author references, and standardizing species names to 'Genus species' format. Additionally, Greengenes2 taxonomies were manually updated to reflect current genus-level designations. GSR-DB contained >500 lineages with descriptions in place of species names (e.g., "marine actinobacterium" and "alpha proteobacterium"); we could not resolve these to valid species names and thus retained these sequences with the existing descriptions. Characteristics and metrics of each database are detailed in Table S1.

#### MARS inoculation of maize and rhizosphere microbiome sample collection

Seeds of *Zea mays* genotype Phz51 were obtained from Thomas Lübberstedt (Iowa State University) (3). Seeds were surface sterilized by soaking in 70% ethanol (10 min) followed by 20% bleach (10 min), with five sterile-water washes after each soak. Seeds were pre-germinated in rolls of germination paper moistened with Murashige and Skoog plant growth medium with macro- and micro-nutrients (4), 1.65 g/L ammonium nitrate, and 2 g/L Captan fungicide. Seedlings were incubated at room temperature for 3 to 5 days until most had a radicle that was 1 cm long.

MARSc is comprised of the model rhizosphere colonist *Pseudomonas putida* KT2440 and 30 bacterial strains isolated from the maize root environment (5–10). MARSc inoculum was prepared by culturing each strain overnight in Reasoner’s 2A (R2A) broth, diluting each culture to an optical density at 600 nm ($OD_{600}$) of 0.1 ± 0.005 in 1 mM phosphate buffer (PB)(0.8 mM potassium phosphate dibasic, 0.2 mM potassium phosphate monobasic, pH 7.1 to 7.4), and mixing equal volumes of the 31 strains.

Pre-germinated seedlings were evenly distributed by size across two treatments (with or without addition of MARSc) and soaked in sterile PB or MARSc inoculum for 1 h at room temperature. After allowing excess inoculum to drain off, seedlings were planted in ethanol-sterilized cones (Ray Leach “Cone-tainer” SC10U) containing 145 g of substrate (75:25 soil:sand mix), using soil from plots fertilized with 120 pounds/acre inorganic fertilizer that were in a long-term variable N fertilizer study site in Boone, Iowa (11). One seedling was placed atop the substrate in each cone and was drenched with 100 µL PB or MARSc inoculum. Cones were then filled to the top with substrate and watered with 10 mL fertilizer (60 mM $NH_4NO_3$, 14.7 mM $KH_2PO_4$, 12.4 mM $K_2SO_4$), corresponding to 120 pounds N fertilizer/acre. Plants were grown for 21 days in a growth chamber with a 16 h/8 h light/dark photoperiod at 24 °C, with substrate moisture maintained at 60-70% water-holding capacity using sterile water and frequent rotation of cone racks to minimize edge effects.

Upon plant removal from cones, loose substrate was removed from the roots using sterile forceps, and a sterile razor blade was used to detach the root at the coleoptile node. Rhizosphere material was separated from the roots by vortexing in sterile PB (10 sec) followed by sonication for 3 min (Branson 5510 ultrasonic cleaner). After sonication, the washate was transferred to a new tube and centrifuged at 7,500 xg for 5 min, the supernatant discarded, and the pellet stored at -80°C. DNA was extracted from pellets using a Qiagen PowerSoil Pro 96-well plate DNA extraction kit, with plates loaded using a modified method to decrease cross-contamination (12).

#### Synthetic spike-in sequences

All amplicon PCR reactions included four bacterial synthetic spike-in sequences (13) – Ec5001, Ga5501, Tb5501, Ec5003 – combined at ratios of 1000:100:10:1, respectively. The spike-in mix was added during barcoding PCR to achieve estimated concentrations spanning 25,000 to 25 copies per 50-µL reaction. The spike-in sequences were manually added to the ACT DB, with the phylum listed as “Bacteria”, class, order, and family listed as “Synthetic”, and genus and species listed as spike-in-specific identifiers. Arbitrary TaxIds were assigned for each rank, ensuring that the four synthetic sequences had identical lineages to the genus level. For unknown reasons, only two of four spike-ins were detected, preventing absolute abundance calculations, so all spike-in reads were later excluded from downstream analyses.

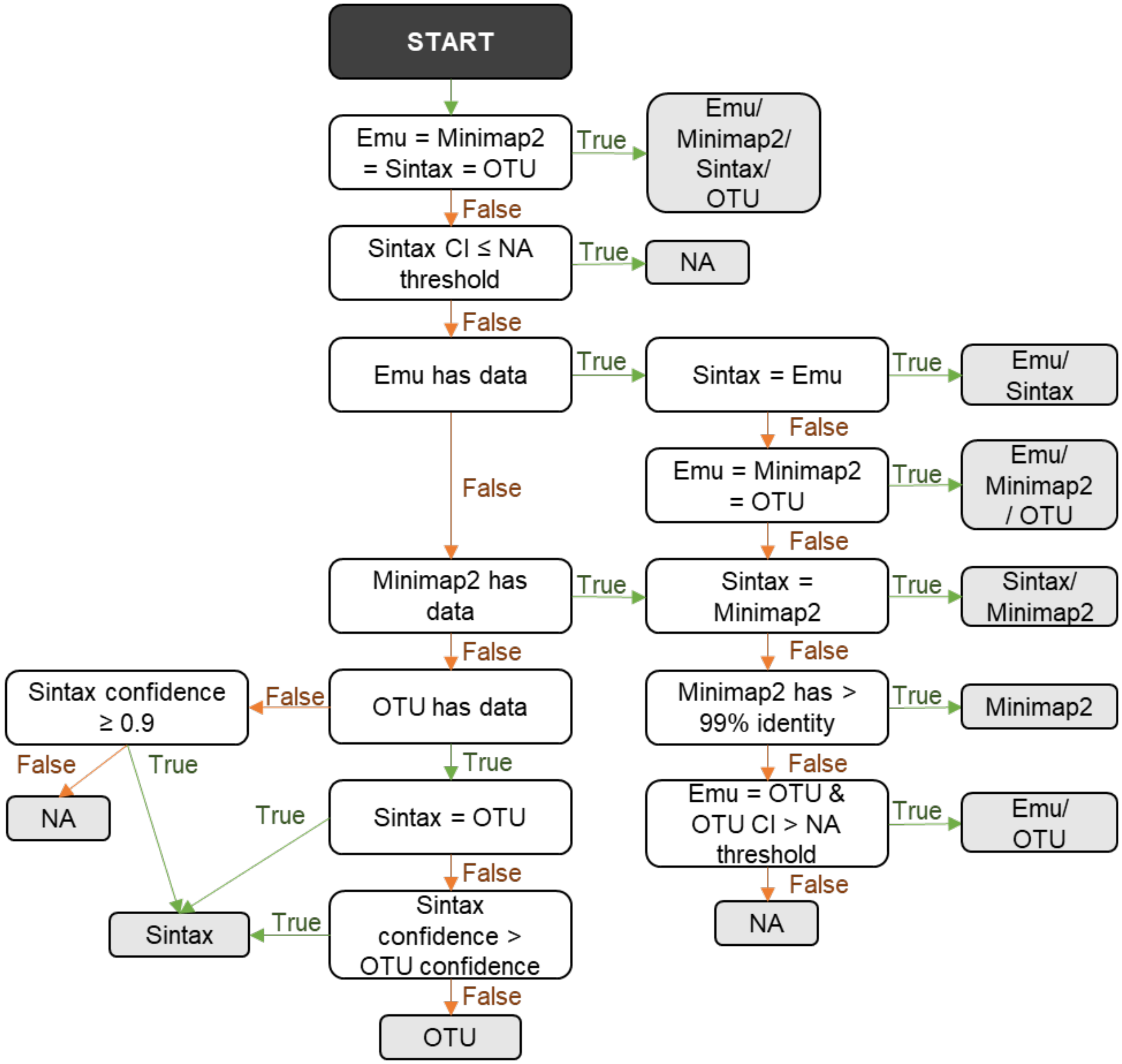


Supplemental Figure 1. ACT decision tree for taxonomic classification of long-read amplicons. The decision tree compares the Sintax, Emu, and LACA outputs on a read-by-read basis for each level of taxonomy. The black node is the beginning and each subsequent white node is a TRUE/FALSE question. Edges are colored and labeled with the corresponding answer. Grey nodes represent the pipeline(s) output selected as the final assignment, e.g., "Emu/Sintax" means that Emu and Sintax agree on the taxonomy and that taxonomy is the one identified as the correct one. "OTU" represents the Sintax taxonomy assignment of the LACA-provided consensus sequence for that OTU. "Minimap2" represents the Minimap2 alignment from EMU before the expectation-maximization polishing. CI, confidence interval. NA, not assigned, where the NA threshold is user defined.

Supplemental Table 1. Characteristics and metrics of the tested reference databases.

| Database | Emu-DB | Emu25-DB | ACT-DB[1] | GSR-DB | GG2[2] | GTDB |
|---|---|---|---|---|---|---|
| **Level of curation** | High | High | High | High | Low | Low |
| **Frequency of updates** | None | N/A | N/A | None | Low | High |
| **Uses NCBI TaxIds** | Yes | Yes | Yes | No | No | No |
| **Supports customization** | No | No | Yes | No | No | No |
| **Number of sequences** | 49,243 | 92,327 | 92,293 | 90,231 | 337,506 | 58,672 |
| **Number of genera represented** | 3,283 | 4,371 | 4,460 | 3,803 | 8,115 | 16,285 |
| **Number of genera with >1 sequence** | 2,067 | 2,868 | 2,985 | 2,577 | 6,092 | 6,630 |
| **Number of species represented** | 17,551 | 27,555 | 17,639 | 18,738 | 22,902 | 58,672 |
| **Number of species with >1 sequence** | 5,879 | 9,984 | 6,609 | 8,178 | 12,586 | 0 |
| **Number of sequences with no genus** | 319 | 56 | 13,477 | 727 | 46,965 | 0 |
| **Number of sequences with no species** | 3 | 1 | 29,555 | 0 | 137,979 | 0 |

[1]Metrics for ACT-DB are specific to the version created for this paper. These numbers may differ depending on the settings used.

[2]GG2, Greengenes2

N/A, not applicable.

Supplemental Table 2. Number of simulated reads for each taxon in the synthetic samples.

| Sample ID[1] | *L. monocytogenes* | *P. aeruginosa* | *B. subtilis* | *E. coli* | *S. enterica* | *L. fermentum* | *E. faecalis* | *S. aureus* | Ec5001[2] |
|---|---|---|---|---|---|---|---|---|---|
| **HL001** | 198,579 | 199,633 | 20,376 | 9,877 | 2,031 | 205 | 20 | 2 | 208 |
| **HL002** | 876,545 | 855,286 | 87,572 | 43,154 | 8,622 | 885 | 85 | 9 | 896 |
| **HL003** | 145,148 | 146,435 | 14,793 | 7,277 | 1,480 | 147 | 14 | 1 | 148 |
| **HL004** | 298,808 | 291,204 | 29,372 | 14,846 | 2,931 | 292 | 29 | 3 | 289 |
| **HL005** | 541,864 | 548,649 | 53,254 | 26,810 | 5,515 | 533 | 54 | 5 | 523 |
| **HL006** | 19,590 | 19,605 | 19,506 | 19,587 | 19,560 | 19,631 | 19,590 | 19,425 | 19,543 |
| **HL007** | 5,656 | 5,605 | 5,626 | 5,625 | 5,663 | 5,708 | 5,687 | 5,659 | 5,677 |
| **HL008** | 8,342 | 8,283 | 8,324 | 8,307 | 8,341 | 8,237 | 8,227 | 8,225 | 8,246 |
| **HL009** | 17,375 | 17,323 | 17,390 | 17,566 | 17,408 | 17,498 | 17,550 | 17,551 | 17,453 |
| **HL010** | 12,867 | 12,735 | 12,661 | 12,759 | 12,702 | 12,822 | 12,904 | 12,817 | 12,831 |
| **HL011** | 19,590 | 19,605 | 19,506 | 19,587 | 19,560 | 19,631 | 19,590 | 19,425 | 0 |
| **HL012** | 5,656 | 5,605 | 5,626 | 5,625 | 5,663 | 5,708 | 5,687 | 5,659 | 0 |
| **HL013** | 8,342 | 8,283 | 8,324 | 8,307 | 8,341 | 8,237 | 8,227 | 8,225 | 0 |
| **HL014** | 17,375 | 17,323 | 17,390 | 17,566 | 17,408 | 17,498 | 17,550 | 17,551 | 0 |
| **HL015** | 12,867 | 12,735 | 12,661 | 12,759 | 12,702 | 12,822 | 12,904 | 12,817 | 0 |

[1]Sample IDs HL001-HL015 were assigned arbitrarily to identify samples in a manner similar to our existing barcode names.

[2]Ec5001 is the synthetic spike in sequence.

Supplemental Table 3. Dataset statistics for samples after read filtering

| Sample name | Library_ Barcode | SRA Accession | Number of seqs | Total bases (mb) | Mean length | N50 | L50 | Q20 (%) | Q30 (%) | Mean Q[1] |
|---|---|---|---|---|---|---|---|---|---|---|
| **Zymo A** | 1_HL067 | SRR37791292 | 37,203 | 56 | 1515.9 | 1513 | 169 | 91.3 | 21.0 | 19.7 |
| **Zymo B** | 2_HL001 | SRR37791291 | 387,290 | 588 | 1518.4 | 1512 | 188 | 91.4 | 20.5 | 19.7 |
| **Zymo C** | 3_HL067 | SRR37791290 | 44,667 | 68 | 1516.2 | 1513 | 178 | 91.5 | 22.2 | 19.8 |
| **Zymo D** | 4_HL001 | SRR37791289 | 278,434 | 422 | 1516.3 | 1512 | 187 | 91.5 | 22.2 | 19.8 |
| **Zymo E** | 1_HL068 | SRR37791296 | 108,482 | 165 | 1516.4 | 1513 | 185 | 91.4 | 20.4 | 19.8 |
| **Zymo F** | 2_HL002 | SRR37791295 | 148,761 | 226 | 1517.2 | 1512 | 182 | 91.4 | 20.6 | 19.8 |
| **Zymo G** | 3_HL068 | SRR37791294 | 109,307 | 166 | 1516.2 | 1512 | 181 | 91.4 | 21.7 | 19.8 |
| **Zymo H** | 4_HL002 | SRR37791293 | 78,785 | 119 | 1516.1 | 1512 | 176 | 91.4 | 22.1 | 19.8 |
| **Rhizo 1** | 4_HL062 | SRR37749542 | 230,068 | 343 | 1492.5 | 1496 | 202 | 90.5 | 20.0 | 19.6 |
| **Rhizo 2** | 4_HL063 | SRR37749541 | 256,990 | 383 | 1492 | 1494 | 205 | 90.6 | 20.6 | 19.6 |
| **Rhizo 3** | 4_HL064 | SRR37749540 | 229,057 | 343 | 1495.7 | 1496 | 204 | 90.6 | 22.3 | 19.6 |
| **Rhizo 4** | 4_HL065 | SRR37749539 | 149,874 | 224 | 1491.6 | 1496 | 205 | 90.7 | 23.3 | 19.6 |
| **Rhizo 5** | 4_HL066 | SRR37749538 | 79,197 | 118 | 1495.5 | 1497 | 192 | 90.5 | 21.7 | 19.5 |

[1]Mean Q is average Phred quality score

Supplemental Table 4. Proportion of ZymoCom reads classified at various taxonomic levels using ACT or Emu with Emu25-DB.

| Pipeline | Taxonomy level | Percent reads classified as Zymo | Percent reads classified as non-Zymo | Percent reads unclassified |
|---|---|---|---|---|
| **ACT** | Domain | 99.95 | 0.00 | 0.04 |
| **ACT** | Phylum | 97.28 | 0.30 | 2.45 |
| **ACT** | Class | 97.10 | 0.44 | 2.45 |
| **ACT** | Order | 94.42 | 0.49 | 5.08 |
| **ACT** | Family | 92.06 | 0.52 | 7.41 |
| **ACT** | Genus | 90.18 | 0.59 | 9.22 |
| **Emu** | Domain | 93.88 | 0.00 | 6.12 |
| **Emu** | Phylum | 93.85 | 0.02 | 6.12 |
| **Emu** | Class | 93.85 | 0.03 | 6.12 |
| **Emu** | Order | 93.84 | 0.03 | 6.12 |
| **Emu** | Family | 93.79 | 0.09 | 6.12 |
| **Emu** | Genus | 93.42 | 0.45 | 6.12 |
